# TbpB-based oral mucosal vaccine provides heterologous protection against Glässer’s disease caused by different serovars of Spanish field isolates of *Glaesserella parasuis*

**DOI:** 10.1101/2024.05.15.594294

**Authors:** Alba González-Fernández, Oscar Mencía-Ares, María José García-Iglesias, Máximo Petrocchi-Rilo, Rubén Miguélez-Pérez, Alberto Perelló Jiménez, Elena Herencia-Lagunar, Vanessa Acebes-Fernández, César B. Gutiérrez-Martín, Sonia Martínez-Martínez

## Abstract

**Background:** *Glaesserella parasuis* has a substantial impact on the pig production as the primary agent of Glässer’s disease, particularly affecting nursery and early fattening stages. Current prophylactic measures, mainly based in serovar-specific bacterins administered parenterally to sows, face limitations due to maternal immunity, which may interfere with the active immunization of piglets. The mucosal administration of TbpB-based subunit vaccines offers a promising approach to overcome these limitations for the control of the disease in weaning piglets. This study evaluates the immunogenicity and heterologous protection of the oral mucosal TbpB^Y167A^ subunit vaccine in colostrum-deprived piglets challenged with four *G. parasuis* clinical isolates belonging to different TbpB clusters and serovars (SVs) recovered from Spanish pig farms.

**Results:** The mucosal administration of a two-dose TbpB-based vaccine induced a robust humoral immune response in immunized colostrum-deprived piglets, significantly increasing IgA (*p <* 0.01) and IgM (*p <* 0.01) concentration 15 days after the second dose. Subsequent infection challenge with four *G. parasuis* clinical isolates demonstrated heterologous protection, markedly improving survival rates (OR: 8.45; CI 95%: 4.97-14.36) and significantly reducing clinical signs and lesions, regardless of the *G. parasuis* TbpB cluster and serovar. The vaccine not only reduced *G. parasuis* colonization in the respiratory tract of immunized piglets (*p* < 0.0001), but also in systemic target tissues, such as the tarsus and carpus joints, liver, and brain (*p* < 0.05). Further immunohistochemical analysis in different lung locations revealed a significantly lower macrophage count in immunized piglets (*p* < 0.0001).

**Conclusions:** Overall, this study demonstrates that the oral mucosal administration of the TbpB^Y167A^ subunits vaccine in piglets provides effective heterologous protection against different virulent European *G. parasuis* field isolates, significantly reducing bacterial colonization and dissemination. These facts position this TbpB-based vaccine as a leading candidate for a universal vaccine against Glässer’s disease.

## Background

*Glaesserella parasuis*, a Gram-negative bacterium belonging to the *Pasteurellaceae* family, is commonly located in the upper respiratory tract of pigs (1). It is the etiological agent of Glässer’s disease in weaning piglets, a syndrome typically characterized by polyarthritis, polyserositis, meningitis and sometimes acute pneumonia, resulting fatal in most cases. This pathogen not only have significant repercussions on mortality and morbidity, but also in key productive parameters, with a substantial economic impact on swine production (2), underscoring the critical need for effective management strategies.

The transmission of *G. parasuis* is predominantly facilitated from sows to piglets (3), with subsequent dissemination among piglets during the nursery phase. While neonatal protection is initially provided by colostrum-deprived maternal antibodies (4), the decline of these antibodies allows the pathogen to potentially overcome innate immunity, infiltrate target tissues, and develop the disease. This challenge is further complicated by the fact that maternal immunity can interfere with active immunization of piglets (5), and the current serovar-specificity of commercially available vaccines (6,7), failing to provide a complete protection in weaning piglets. The absence of a multivalent vaccine that can overcome maternal immunity contributes to the persistence of Glässer’s disease in pig production (8), urging for the development of innovative approaches that may circumvent these limitations.

Subunit vaccines, which contain antigenic molecules specific to this microorganism may be useful, but they are not currently commercially available. Transferring-binding proteins (Tbp), which confer *G. parasuis* its ability to acquire iron from porcine transferrin to support its growth in low-iron environments, such as mucosal surfaces (9), have been proposed as vaccine antigens (10). A mutated form of TbpB, TbpB^Y167A^, originating from *G. parasuis* Nagasaki strain - TbpB cluster III, serovar 5 (SV5) - and characterized by its impaired ability to bind porcine transferrin, has been demonstrated to display a drastic reduction in its binding affinity (11). Besides, its mucosal administration may avoid the interference of maternal immunity (12), constituting a good candidate for the control of Glässer’s disease in weaning piglets.

The aim of this study is to evaluate the immunogenicity and heterologous protection of the oral mucosal TbpB^Y167A^ subunit vaccine in colostrum-deprived piglets challenged with four *G. parasuis* clinical isolates belonging to different SVs and TbpB clusters recovered from Spanish pig farms.

## Methods

### Selection of *Glaesserella parasuis* field isolates for infection challenge and inoculum preparation

Following their serotyping and phylogenetic analysis (13), four clinical field isolates of *G. parasuis* recovered from Spanish pig farms were selected to evaluate the efficacy of a mucosal TbpB^Y167A^ subunit vaccine against Glässer’s disease in weaning piglets (**Table 1**).

**Table 1.** Selection of *Glaesserella parasuis* field isolates used for infection challenge.

| Isolate ID | Pathologic process | Serovar | TbpB cluster |
| --- | --- | --- | --- |
| Gp-12 | Systemic | 5 | III |
| Gp-39 | Respiratory | 4 | I |
| Gp-46 | Systemic | 1 | I |
| Gp-52 | Respiratory | 7 | III |

For infection challenge, the selected field isolates were initially cultured on chocolate agar plates (Oxoid, UK) at 37 °C for 48 hours. Subsequently, the isolates were incubated for 12 hours in 10 ml of pleuropneumonia-like organisms (PPLO) broth (Pronadisa, Spain) supplemented with glucose (5 µg/ml) and nicotinamide adenine dinucleotide (NAD) (Sigma-Aldrich, USA) (2 µg/ml). A total volume of 1 ml of the cultured broth was transferred to a new flask with 100 ml of PPLO broth until reaching a concentration of 1×10^7^ CFU/ml. All *G. parasuis* incubations were performed under microaerophilic conditions. Bacterial suspensions were further centrifuged at 3,000 rpm during 30 minutes at 6 °C and resuspended in the same volume of RPMI 1640 broth (Gibco, USA) for intranasal piglet infection. A 500 µl total dose was administered per piglet (250 µl per nostril).

### Antigen preparation and vaccine formulation

Vaccine antigen was generated from the binding-defective TbpB^Y167A^ mutant following established protocols (11,14).Vaccine formulations used in immunization studies were prepared following previously documented protocols (12). In summary, they were produced in sets of 50 doses. Each 50-dose composition contained 1.25 mg of antigen (equivalent to 25 µg of protein antigen per dose), 50 µg of poly I:C (Invivogen, USA), and 1 ml (20% v/v) of Montanide Gel 01 adjuvant (Seppic, France). The formulations were adjusted to a final volume of 5 ml using sterile Dulbecco’s Phosphate-Buffered Saline (DPBS) (Sigma-Aldrich, USA). Vaccine formulations underwent mild agitation employing a tabletop rotary shaker at room temperature overnight, followed by subsequent storage at 4 °C until use.

### Experimental design and ethical considerations

This study involved 38 Landrace x Large-White colostrum-deprived piglets, reared artificially according to the protocol described by Guizzo *et al.* (14). Prior to the experiment, all piglets were confirmed free of *G. parasuis* via PCR analysis of nasal swabs. The piglets were randomly allocated into four groups, each comprising 9 to 10 animals, and housed in separate rooms within a biosafety level 2 facility.

The experiment was conducted in two phases. In the initial phase, five piglets from each group received oral mucosal immunization with two doses (administered on days 15 and 30) of the TbpB^Y167A^ mutant vaccine using the *Comfort-in* needle-free device (Gamastech, Italy). The rest of the piglets served as non-immunized controls and were inoculated twice with equal volume of phosphate-buffered saline (PBS). The second phase began on day 45, 15 days post-administration of the second vaccine dose. Each group of piglets was challenged intranasally with 500 µl (250 µl per nostril) of a unique *G. parasuis* field isolate, with a final dose of 5×10^6^ CFU. This challenge phase lasted 7 days, after which all remaining piglets were euthanized with an intravenous overdose of sodium pentobarbital (LeonVet, Spain). Any animal exhibiting severe distress throughout the infection challenge was humanely euthanized before the end of the experiment. Figure 1 presents a detailed scheme of the experimental design.

**Figure 1.**
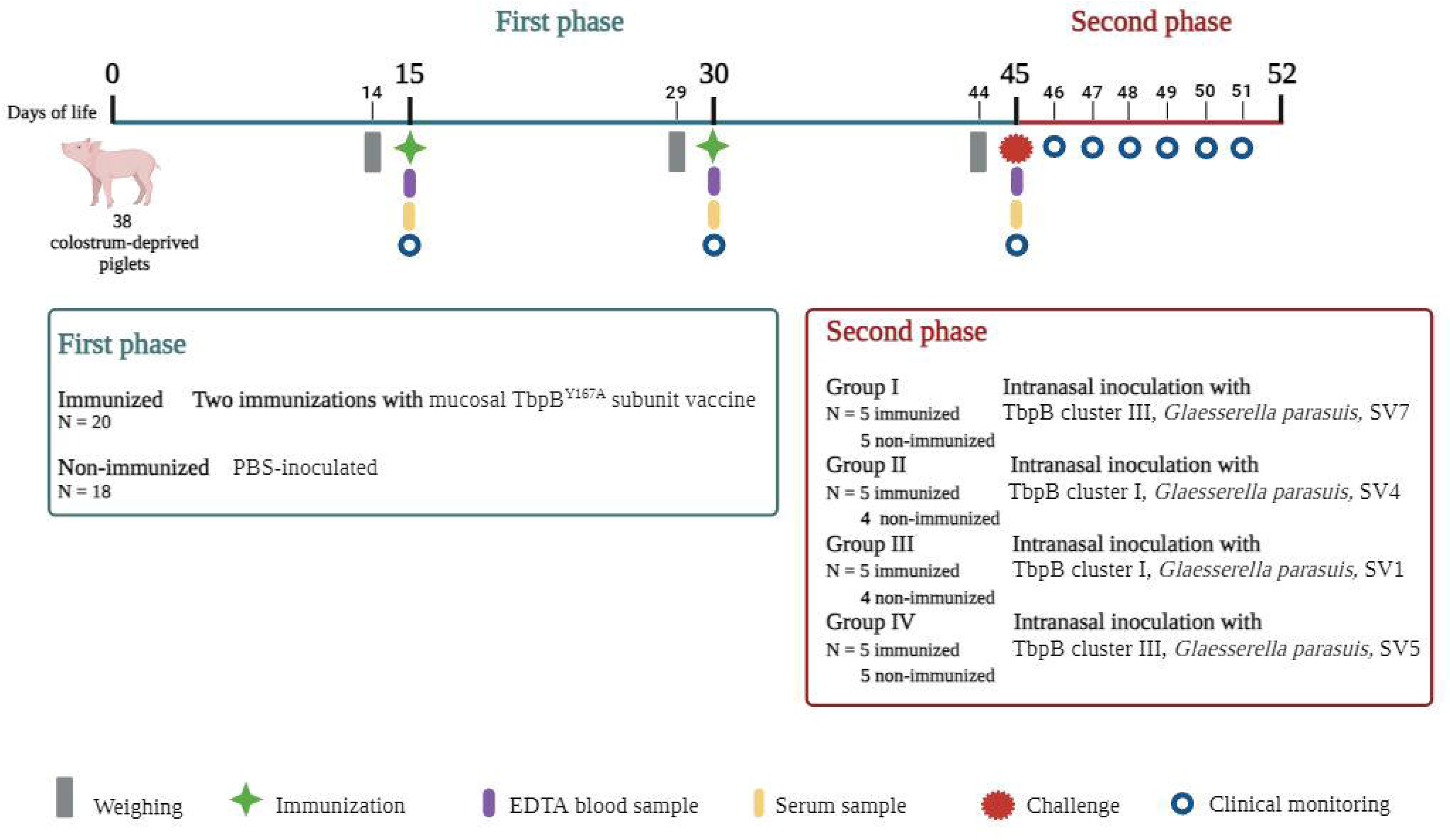
Two-phase experimental design for the evaluation of the immunogenicity and efficacy of the TbpB-based oral mucosal vaccine against Glässer’s disease. The first phase (marked in blue) consisted of the immunization of colostrum deprived piglets. Immunized piglets (*n* =20) received two oral mucosal administrations of the TbpB^Y167A^ subunit vaccine, while non-immunized piglets (*n* = 18) were inoculated with PBS. The second phase (marked in red) began 15 days after the second dose of the vaccine and piglets were divided into four groups and intranasally challenged with four different *Glaesserella parasuis* field isolates. This challenge lasted for seven days, after which all remaining piglets were euthanized. Animals were daily monitored for clinical signs and temperature and those exhibiting severe distress were humanely euthanized before the end of the experiment. After euthanasia, a necropsy was performed to evaluate macroscopic lesions and collect samples. Serum samples were collected on days 15, 30 and 45. SV: serovar.

All procedures involving the piglets were approved by the University of León Committee on Animal Care and Use (OEBA-ULE-003-2022). Veterinary professionals and trained personnel, complying with both Spanish and European Union regulations, conducted the handling of the animals. Animals were clinically examined upon arrival and monitored throughout the experiments.

### Clinical monitoring and sample collection during challenge

During the initial phase, serum samples were collected on three occasions: at the time of the first vaccine dose (day 15), the second dose (day 30), and the beginning of the second phase (day 45). Post-collection, the serum underwent centrifugation at 2,500 rpm for 10 minutes at 8 °C following a 3-hour incubation at 4 °C. Aliquots of the resultant serum were stored at −80 °C until further use. In this phase, an exhaustive clinical follow-up was carried out to evaluate possible adverse reactions to the vaccine.

In the second phase, daily monitoring of rectal temperature and clinical signs was conducted. Fever was considered above 40 °C. Clinical signs were individually scored (0 = no clinical signs; 1 = mild clinical signs; 2 = moderate clinical signs; and 3 = severe clinical signs) across six categories: apathy, lack of appetite, lameness, incoordination, dyspnea, and joint inflammation. Animals presenting moderate to severe clinical signs were humanely euthanized to prevent suffering, followed by necropsy. Macroscopic examination was performed paying special attention to the joints, serous (peritoneum, pericardium, and pleura), and to different organs, as spleen, liver, kidneys, and brain. The macroscopic lesions of each experimental group were photographed for comparison. Inflammatory and vascular lesions (edema, congestion, hemorrhages and fibrin deposit) were scored using the same 0-3 scale described above.

Swabs for the evaluation of the growth of *G. parasuis* were collected from tarsus, carpus and atlantooccipital joints, abdominal cavity, pericardium, lungs, spleen, liver and brain. Then, they were cultured on chocolate agar plates at 37 °C for 48 hours under microaerophilic conditions. The identity of the bacterium was validated through matrix-assisted laser desorption-ionization time-of-flight mass spectrometry (MALDI-TOF MS) (Bruker, Germany). Tissue samples were collected from liver, gallbladder, spleen, lungs, heart, and brain for histological and immunohistochemical (IHC) studies. Bronchoalveolar lavage fluid (BALF) was also collected by introducing sterile PBS using a 20 ml syringe through the trachea and ensuring that the solution reached the alveoli by rubbing. Subsequently, the lungs were inverted to introduce the resulting fluid into a sterile 50 ml tube. Afterwards, the tubes were centrifuged twice at 2,000 rpm for 5 minutes and the supernatant was collected in aliquots and further frozen at −80 °C until use.

### Evaluation of the humoral immune response

IgM and IgA levels in serum and BALF, respectively, were quantified using ELISA commercial kits (Antibodies, UK), following the manufacturer’s protocols. IgG levels against *G. parasuis* and TbpB^Y167A^ antigen were quantified on serum samples using two in-house ELISA based on the methodology described by Martín de la Fuente *et al.* (10). Briefly, ELISA plates were coated by the addition of 100 µl of either complete inactivated *G. parasuis* or TbpB^Y167A^ antigen diluted in carbonate buffer (5 µg per well). This was followed by a 2-hour incubation at 37 °C and then an overnight incubation at 4 °C. Plates were subsequently washed three times with 300 µl/well of PBS containing 0.05% Tween 20 (PBS-T) (Sigma-Aldrich, USA). Non-specific binding sites were blocked using 300 µl/well of 3% (w/v) skim milk powder (Sigma-Aldrich, USA) in PBS-T. Plates were then washed three times with 300 µl/well of PBS-T. After coating, 100 µl of diluted porcine serum (1:100) was added to each well and incubated for 60 minutes at 37 °C. Plates were then washed three times with 300 µl/well of PBS-T. Following this, 100 µl/well of peroxidase-conjugated goat anti-swine IgG (Sigma-Aldrich, USA) was then applied, followed by another 60-minute incubation at 37 °C. Plates were subsequently washed three times with 300 µl/well of PBS-T. This was succeeded by the addition of 100 µl/well TMB substrate (Sigma-Aldrich, USA) containing 0.002% H_2_O_2_ for 10-15 minutes. The reaction was terminated with 100 µl/well of 3 M HCl, and absorbance was measured at 450 nm. Immunoglobulin levels were quantified in duplicate and expressed in mg/ml, using a calibration curve.

### Histological and immunohistochemical assays

Tissue samples obtained from the necropsies were fixed in 10% buffered formalin. Subsequently, they were embedded in paraffin and cut into 3-4 µm sections using a microtome for subsequent staining with hematoxylin-eosin (HE). Microscopic lesions were individually scored (0 = no lesion; 1 = minor lesion; 2 = mild lesion; 3 = moderate lesion; and 4 = severe lesion).

To perform the IHC assay, lung samples previously embedded in paraffin were selected. They were cut into 3 µm thick sections, placed on poly-L-lysine-treated slides (Menzel Gläser, Germany) and allowed to dry at room temperature. The samples were kept overnight in an oven at 60 °C to facilitate deparaffinization. Slides were deparaffinized using xylene (VWR, Spain) and gradually rehydrated with a series of alcohol gradients until reaching distilled water. Subsequently, the slides underwent heat-mediated antigen retrieval treatment, following the procedure outlined by Domenech *et al.* (15), with modifications. Briefly, the samples were immersed in 0.01 M citrate buffer, pH 6.0, at 90 °C for 40 minutes. The tissue sections rested for 20 minutes at room temperature and then they were immersed in distilled water for 5 minutes, followed by two washes with PBS-T buffer, pH 7.6, for 5 minutes each. To block nonspecific reactions, the samples were covered with 150 µl of horse serum diluted 1/20 in PBS, 0.1% bovine serum albumin (BSA) (Sigma-Aldrich, USA), and sodium azide (Sigma-Aldrich, USA) for 25 minutes in a humid chamber. After removing the excess serum from the tissue sections, 150 µl of the hybridoma supernatant CD163 (mouse monoclonal antibody, clone 2A10/11) (INIA, Spain) was added overnight at 4°C in a humid chamber. After two washes with PBS-T, 150 µl of a 1/200 dilution biotinylated anti-mouse antibody (Vector Laboratories, USA) was added for 30 minutes at room temperature in a humid chamber. Endogenous peroxidase was inhibited by distilled water with 0.5% H_2_O_2_ in darkness for 30 minutes after two washes with PBS-T. Two additional washes with PBS-T for 5 minutes each were performed, and tissue sections were covered with 150 μL of Avidin-Biotin-Peroxidase (ABC) Elite complex (Vector Laboratories, USA) for 40 minutes in a humid chamber at room temperature, followed by two washes with PBS-T. The sections were then covered with 150 µl of NovaRed chromogen (Vector Laboratories, USA), and the development was monitored by optical microscope visualization every 3-4 minutes until the nuclei began to show a reddish color. To assess nonspecific immunostaining in tissue sections, negative controls were included, which were not incubated with the hybridoma supernatant CD163. Instead, they were solely covered with the diluent PBS, 0.1% BSA and sodium azide. Slides were counterstained with Harris hematoxylin.

The immunoreaction was evaluated using light microscopy (Eclipse E600, Nikon, Japan) with a Nikon Digital Sight DS-Fi1® camera (Nikon, Japan). Areas with higher positivity (*hot spots*) were selected at low magnifications (objective of x4 and x10) for the quantitative analysis of the immunostaining. Ten non-overlapping areas from alveoli, bronchi, bronchioles, blood vessels, and interstitium were selected and macrophages were counted at 400x magnification (area 0.035 mm² x 10 fields = 0.35 mm²) by the manual count tool of the NIS-Elements BR software (Nikon Instruments Inc., Japan). Then, the arithmetic mean and the standard deviation were calculated.

### Statistical analyses and results visualization

The evaluation of vaccine immunogenicity before the infection challenge, including levels of IgA, IgM, IgG against *G. parasuis*, and IgG against TbpB^Y167A^, was conducted using non-parametric statistical methods due to the non-normal distribution of the data. Specifically, differences in immunoglobulin levels between immunized and non-immunized groups were assessed using the Wilcoxon rank-sum test. To examine the temporal changes in immunoglobulin levels within the immunized group, the Wilcoxon signed-rank test, a test for paired data, was employed.

For the analysis of infection challenge outcomes, metadata including piglet identification, immunization status (immunized or non-immunized), TbpB cluster (I or III), and the SV (1, 4, 5, or 7) of the *G. parasuis* isolate were integrated. Key variables analyzed encompassed clinical scores, rectal temperature, survival rates, bacterial growth in various tissues, the scoring of macroscopic and microscopic lesions and macrophage counts in lung sections via IHC. Statistical comparisons were initially made between immunization groups and then between immunized and non-immunized piglets within each specific TbpB cluster and SV of *G. parasuis* involved in the challenge. All statistical analyses were conducted using the Wilcoxon rank-sum test. For the evaluation of the survival rate, an additional Cox Model was performed to evaluate the protective effect of the vaccine for the evaluated groups, providing the information expressed as odds ratio (OR) and confidence interval 95% (CI 95%).

All analyses were conducted using R version 4.3.2 (2023-10-31 ucrt) (16). *P*-values were adjusted by following the Benjamini & Hochberg method (17) and significance was established at *p* < 0.05. Plots were produced using the *ggplot2* package (18) and further modified using the software *Inkscape* version 1.3.2 (https://inkscape.org/). The level of statistical significance was represented with asterisks: four asterisks (****) indicated a *p*-value less than 0.0001; three asterisks (***) indicated a *p-*value between 0.0001 and 0.001; two asterisks (**) indicated a *p-*value between 0.001 and 0.01; one asterisk (*) indicated a *p-*value between 0.01 and 0.05; non-significance (ns) indicated a *p-*value higher than 0.05.

## Results

### Evaluation of the humoral immune response prior to infection challenge

When comparing the humoral immune response in immunized and non-immunized colostrum-deprived piglets, we could observe a significant increase (*p* < 0.01) of all immunoglobulins in immunized animals 15 days after the first (day 30) and second dose (day 45) of TbpB^Y167A^ mucosal subunit vaccine, except for IgG against TbpB^Y167A^ at day 30 **(Figure 2).** On immunized animals, a significant upward trend (*p* < 0.05) in all evaluated immunoglobulins was demonstrated on day 30 and, particularly, on day 45. It is particularly remarkable the noteworthy increase in IgM and IgA concentration when compared to IgG levels (*p* < 0.01), suggesting an increased mucosal protection against *G. parasuis*. A detailed information of the mean and standard deviation of each immunoglobulin along the study is available in an additional file (see Additional file 1).

**Figure 2.**
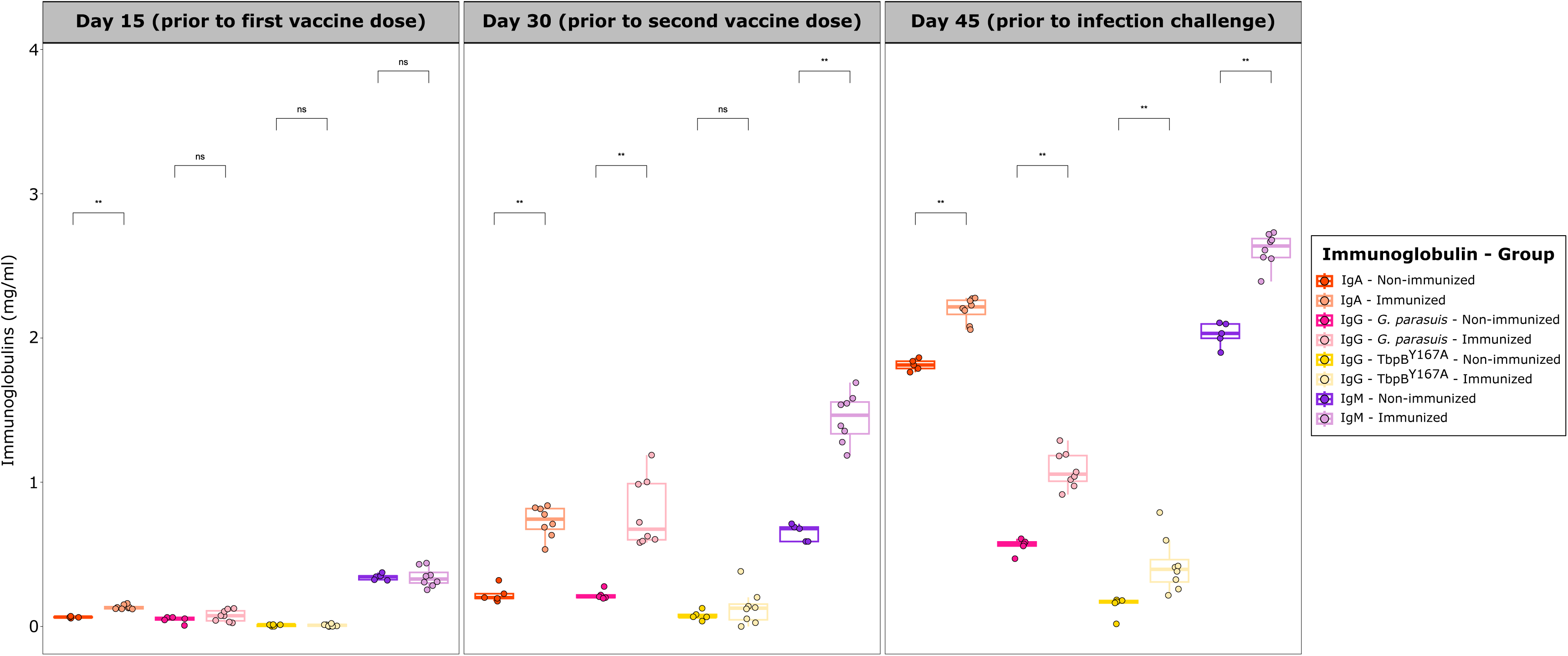
Evolution of immunoglobulin concentration (mg/ml) in immunized and non-immunized colostrum-deprived piglets after oral mucosal administration of the two-dose TbpB-based vaccine. Each piglet is represented by a dot with horizontal jitter in the boxplot for visibility. The horizontal box lines represent the first quartile, the median, and the third quartile. Whiskers include the range of points within the 1.5 interquartile range. Differences per group were evaluated with the Wilcoxon rank-sum test.

### Efficacy of the immunization to prevent mortality and to reduce morbidity after *G. parasuis* infection

Following the infection challenge with *G. parasuis* field isolates, a highly significant protection was conferred by the vaccination in the survival rate (*p* < 0.0001), regardless of *G. parasuis* TbpB cluster and SV, since all non-immunized piglets died within the initial three days post-infection (dpi). Indeed, the vaccine increased by eight times the survival chance of animals challenged with the pathogen (OR: 8.45; CI 95%: 4.97-14.36). This protective effect was particularly high for animals challenged with SV5 (OR: 9.59; CI 95%: 3.34-27.50) and, especially, with SV4, for which no OR could be established due to the extremely high differences between groups (*p <* 1×10^-9^), with all immunized animals surviving the whole study (Figure 3A). A detailed description of the protective effect of vaccination among groups is available in an additional file (see Additional file 2).

**Figure 3.**
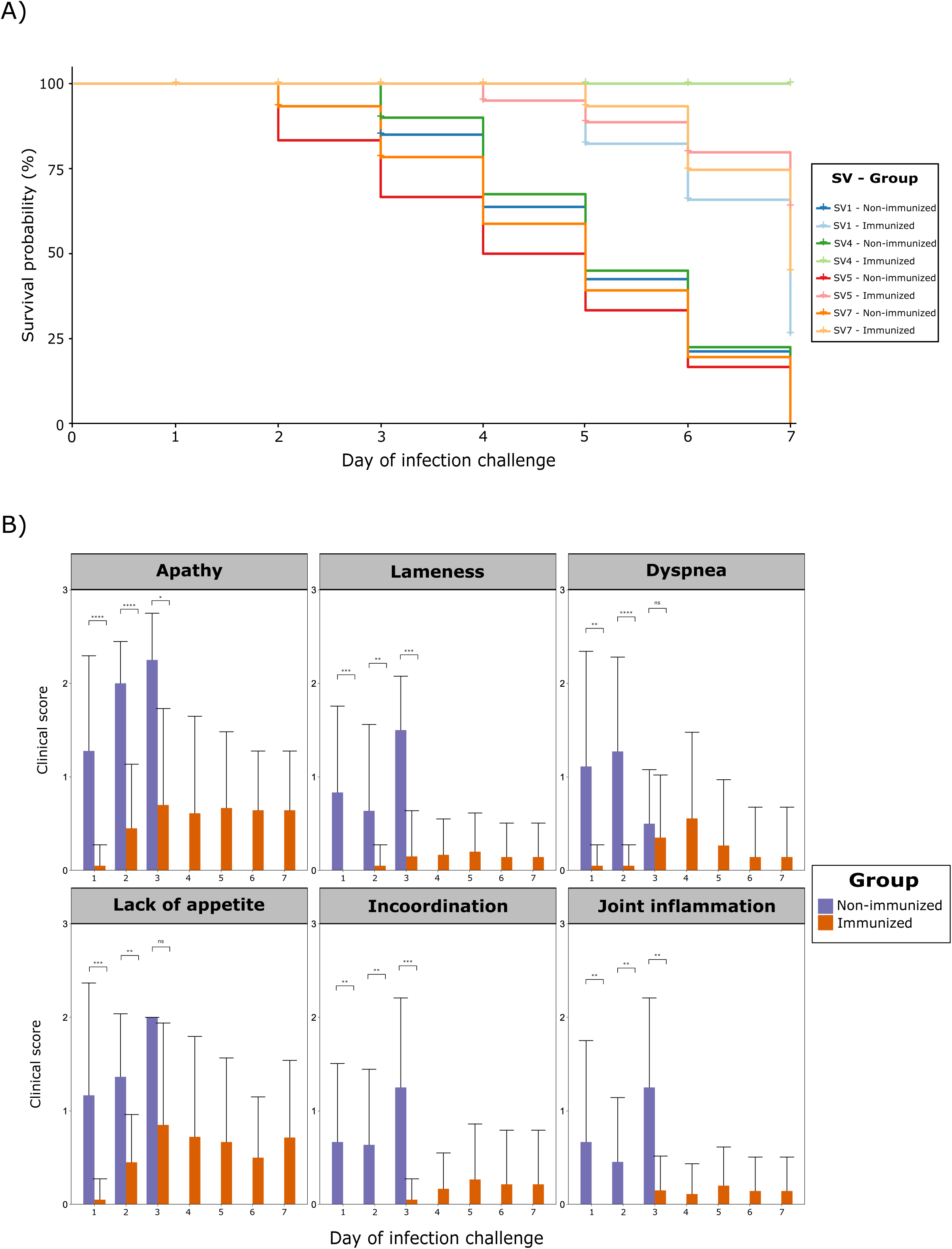
Evaluation of the efficacy of the oral mucosal TbpB-based vaccine in piglets immunized against Glässer’s disease throughout a seven-day experiment. A) Survival rate, expressed as percentage, of immunized and non-immunized piglets challenged with four different serovars (SVs) of *Glaesserella parasuis*. B) Clinical score average (0–3) and standard deviation of immunized and non-immunized piglets, itemized into six different clinical signs. No non-immunized piglet survived after the third day of challenge. Differences per group were evaluated with the Wilcoxon rank-sum test.

Daily monitoring of rectal temperature unveiled a significantly higher temperature among non-immunized piglets (*p* < 0.05), regardless of TbpB cluster or SV, which was particularly pronounced during the first dpi. Further analysis within *G. parasuis* TbpB clusters revealed differential responses to vaccination. Notably, this temperature was higher in non-immunized animals challenged with *G. parasuis* isolates belonging to cluster III on day 1 post-infection (*p* < 0.05), and on cluster I on days 2 and 3 (*p* < 0.05), underscoring the impact of vaccination on temperature dynamics across these clusters. Additionally, within animals challenged with *G. parasuis* SV7, a constantly higher rectal temperature was observed in non-immunized piglets (*p* < 0.05).

Clinical monitoring revealed that all non-immunized piglets exhibited clinical signs from the first dpi, with consistently higher scores among non-immunized animals across different dpi, demonstrating the protective effect of the vaccine at the symptomatic level (Figure 3B). This finding was consistent when evaluating the different *G. parasuis* TbpB clusters, but remarkable differences were observed among SVs, with a higher clinical score among non-immunized piglets. For instance, dyspnea predominated on the second dpi for SV1(*p* < 0.05), lameness on the third dpi for SV4 (*p* < 0.05), incoordination on the second dpi for SV7 (*p* < 0.05), and a combination of apathy, dyspnea, inflammation, and lameness on the first dpi for SV5 (*p* < 0.05). A detailed description of differences for rectal temperature and clinical signs between immunized and non-immunized piglets, itemized by TbpB cluster and SV, is available in an additional file (see Additional file 3).

### Evaluation of the vaccine protection based on macroscopic and microscopic lesions after *Glaesserella parasuis* infection

When considering macroscopic lesions, a consistently higher lesions score was observed in organs from non-immunized piglets, demonstrating the protective effect of the vaccine This effect was most evident in TbpB cluster III animals in which lesions were consistently greater in non-immunized animals challenged with SV5 and SV7 compared to those challenged with SV1 and SV4 (TbpB cluster I) (*p* < 0.01) (Figure 4). Notably, gallbladder edema and pleuritis exhibited the larger differences (*p* < 0.001). It is remarkable that immunized animals challenged with *G. parasuis* TbpB cluster III exhibited significantly lower score for pericarditis (*p* < 0.01) (Figure 4C and D) and pleuritis (*p* < 0.001) (Figure 4E and F) than non-immunized animals. This finding was also observed in peritonitis, although significant differences were only found in SV5 (*p* < 0.05) and not in SV7 (Figure 4A and B). Several differences were also observed within each SV, highlighting, the protective effect of vaccination in piglets challenged with SV5, revealing a significantly lower score for arthritis (*p* < 0.01), peritonitis (*p* < 0.05), pericarditis (*p* < 0.05), and pleuritis (*p* < 0.05) in immunized piglets than non-immunized ones in comparison to the other SVs.

**Figure 4.**
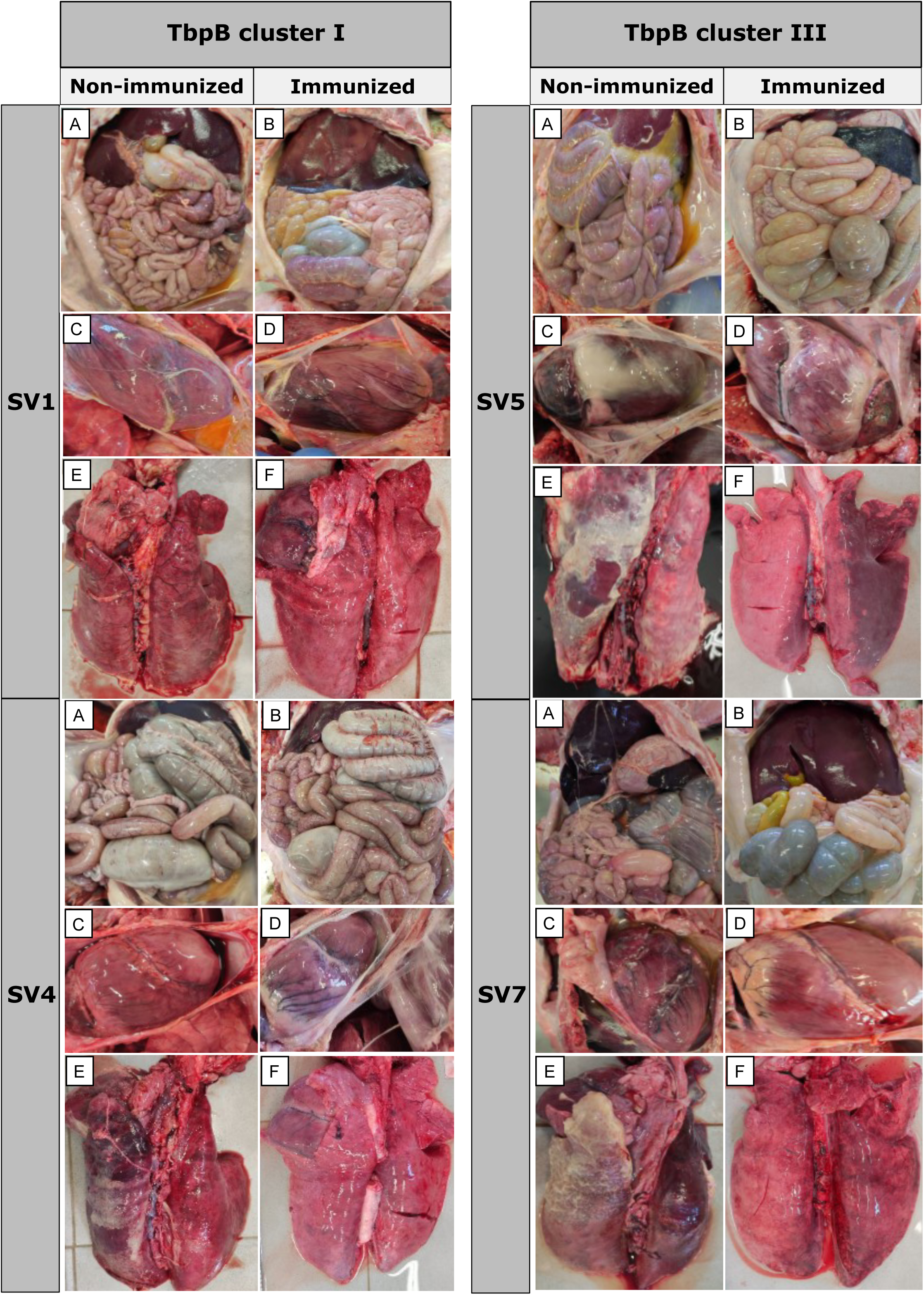
Comparison of macroscopic lesions after *Glaesserella parasuis* infection challenge between immunized and non-immunized piglets with the oral mucosal TbpB-based vaccine. Abdominal and thoracic cavities are shown and itemized by *G. parasuis* TbpB cluster and serovar (SV). A and B, peritoneal serosae; C and D, pericardium; E and F, pleura.

Histopathological examination conducted upon completion of the experiment demonstrated that, surprisingly, the vascular lesion scores were significantly higher in immunized animals when compared to non-immunized ones (*p <* 0.05), highlighting the alveolar and/or interstitial edema (*p* < 0.001), liver congestion (*p* < 0.05) and gallbladder edema (*p* < 0.05), with no differences in spleen or brain lesions. These differences were particularly high in animals challenged with heterologous *G. parasuis* TbpB cluster I, with a significantly higher scores for lesions in immunized animals (*p* < 0.05), not only in the lesions mentioned above, but also in brain ventriculitis (*p* < 0.05). It is remarkable that no differences were observed for animals challenged with *G. parasuis* TbpB cluster III. Within TbpB cluster I, SV1 showed moderate to severe meningoencephalitis in the immunized piglets in contrast to the absence of this inflammation in the non-immunized ones, which had survived only 2-3 days after challenge (Figure 5A and B). Interestingly, the only score lesion that was significantly lower in immunized animals was the interstitial pneumonia for animals challenged with *G. parasuis* SV5 (*p* < 0.01) (Figure 5 C and D), revealing the vaccine protection against inflammatory lung disorders in immunized piglets. A detailed description of differences for macroscopic and microscopic lesions between immunized and non-immunized piglets, itemized by TbpB cluster and SV, are available in an additional file (see Additional file 4).

**Figure 5.**
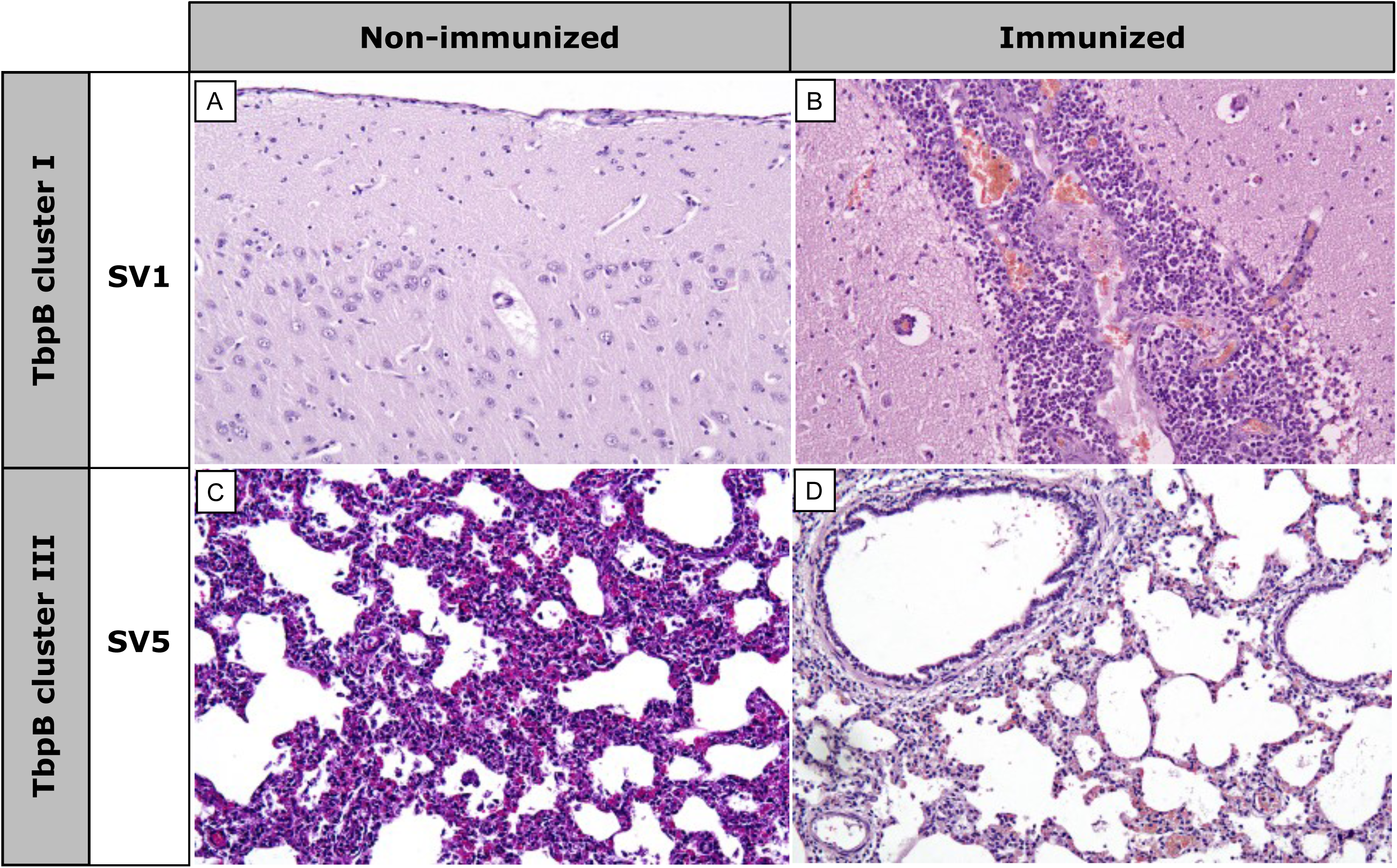
Histopathological differences after *Glaesserella parasuis* infection challenge between immunized and non-immunized piglets with the oral mucosal TbpB-based vaccine. Brain (A and B) and lung (C and D) are itemized by *G. parasuis* TbpB cluster and serovar (SV). Non-immunized piglets infected with TbpB cluster I (SV1) showed no brain lesions (A), while immunized ones revealed intense meningitis (B). Non-immunized animals challenged with TbpB cluster III (SV5) showed interstitial pneumonia (C), while absence or slight pneumonia was observed in immunized ones (D). X100.

### *Glaesserella parasuis* colonization in target tissues of piglets after infection challenge

*G. parasuis* growth on samples recovered from different tissues after necropsy revealed a significantly higher presence of the pathogen in non-immunized animals (*p* < 0.05), except for abdominal cavity and the atlantooccipital joint (Figure 6). It is particularly remarkable that 94% of lungs from non-immunized piglets were positive to *G. parasuis* growth, while barely 15% of immunized were positive to this pathogen (*p* < 0.0001). Upon stratifying by *G. parasuis* TbpB cluster, microbial growth in lungs was consistent with the previous finding in both clusters (*p* < 0.01), while notable distinctions emerged within each cluster. For instance, non-immunized animals challenged with *G. parasuis* cluster I exhibited a significantly higher growth in carpus, tarsus joints (*p* < 0.05), whereas those challenged with cluster III had a higher presence of *G. parasuis* in the liver (*p* < 0.05). Interestingly, when itemizing by SVs, its growth in lungs was consistent for SV1 (*p* < 0.05), SV4 (*p* < 0.05) and SV7 (*p* < 0.001), but not for SV5. We also observed that the vaccine reduced the growth of *G. parasuis* in carpus and tarsus joints of piglets challenged with SV4. All these findings underscore the diverse impact of vaccination across anatomical sites and the complex interplay between vaccine response and bacterial colonization dynamics. A detailed information of bacterial growth between immunized and non-immunized piglets, itemized by TbpB cluster and SV, is available in an additional file (see Additional file 5).

**Figure 6.**
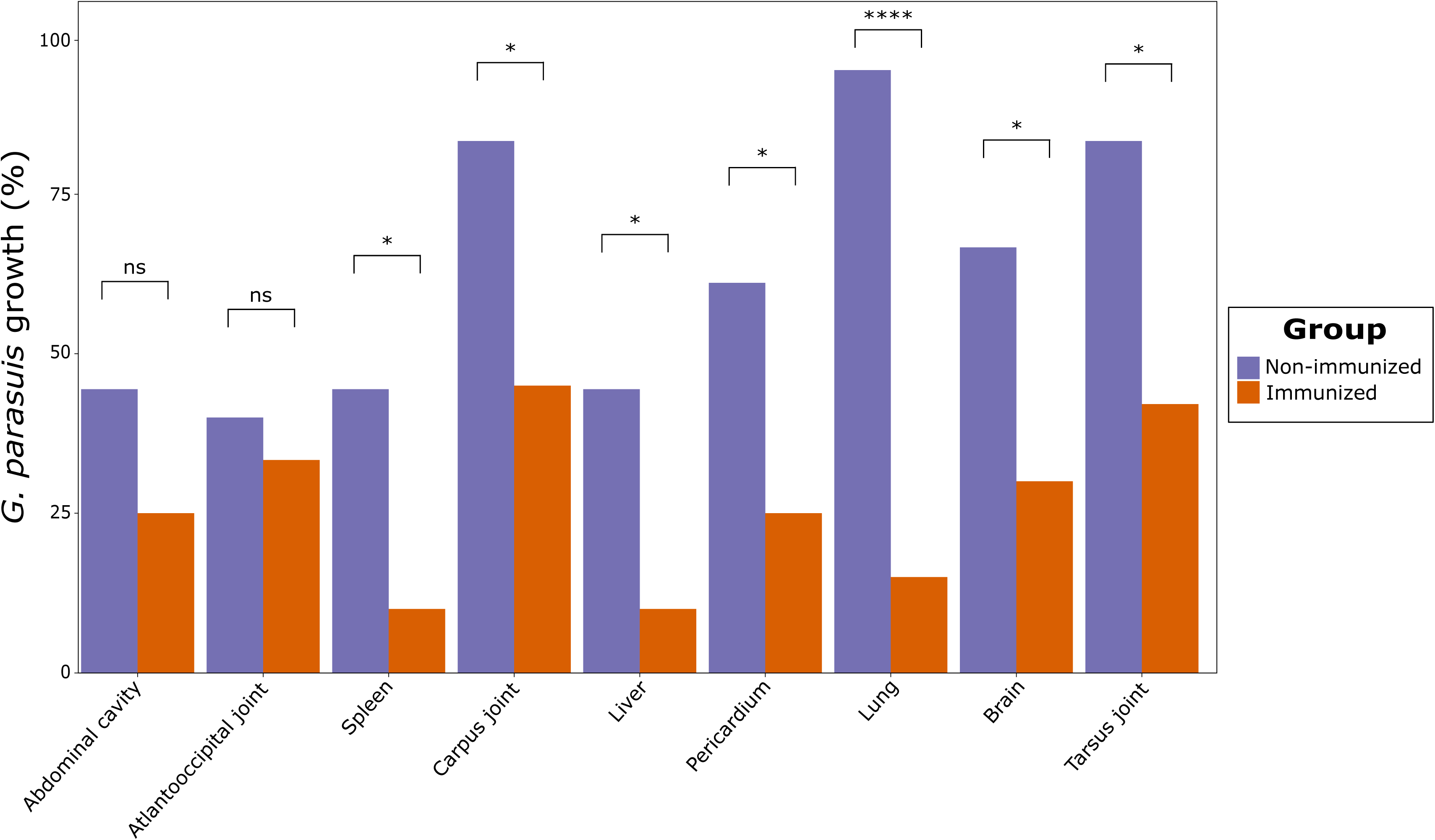
*Glaesserella parasuis* growth, expressed as percentage, in different anatomical regions after infection challenge in immunized and non-immunized piglets with the oral mucosal TbpB-based vaccine. Differences per group were evaluated with the Wilcoxon rank-sum test.

### Determination of macrophage abundance in pulmonary tissue

IHC quantification of labeled macrophages across various lung locations revealed a consistently high number in non-immunized animals in all evaluated lung sections (*p* < 0.01), demonstrating the protective effect of the vaccine (Figure 7). These differences were consistent for animals challenged with *G. parasuis* TbpB cluster I in all lung sections (*p* < 0.01), while these differences were not observed in bronchi and bronchiole in cluster III. When itemizing by SVs, we observed remarkable differences, with a persistent significantly higher number of macrophages in lung lesions in non-immunized animals challenged with SV1 (*p* < 0.05) and SV4 (*p* < 0.05), or the higher presence of macrophages in the interstitium and alveoli in non-immunized piglets challenged with SV5 and SV7 (*p* < 0.05). Altogether, a lower total recount of macrophages was observed in immunized piglets (*p* < 0.05), regardless of the *G. parasuis* SV, which reveals the protective effect of the vaccine across all SVs. A detailed comparison of macrophage abundance between immunized and non-immunized piglets, itemized by TbpB cluster and SV, is available in an additional file (see Additional file 6).

**Figure 7.**
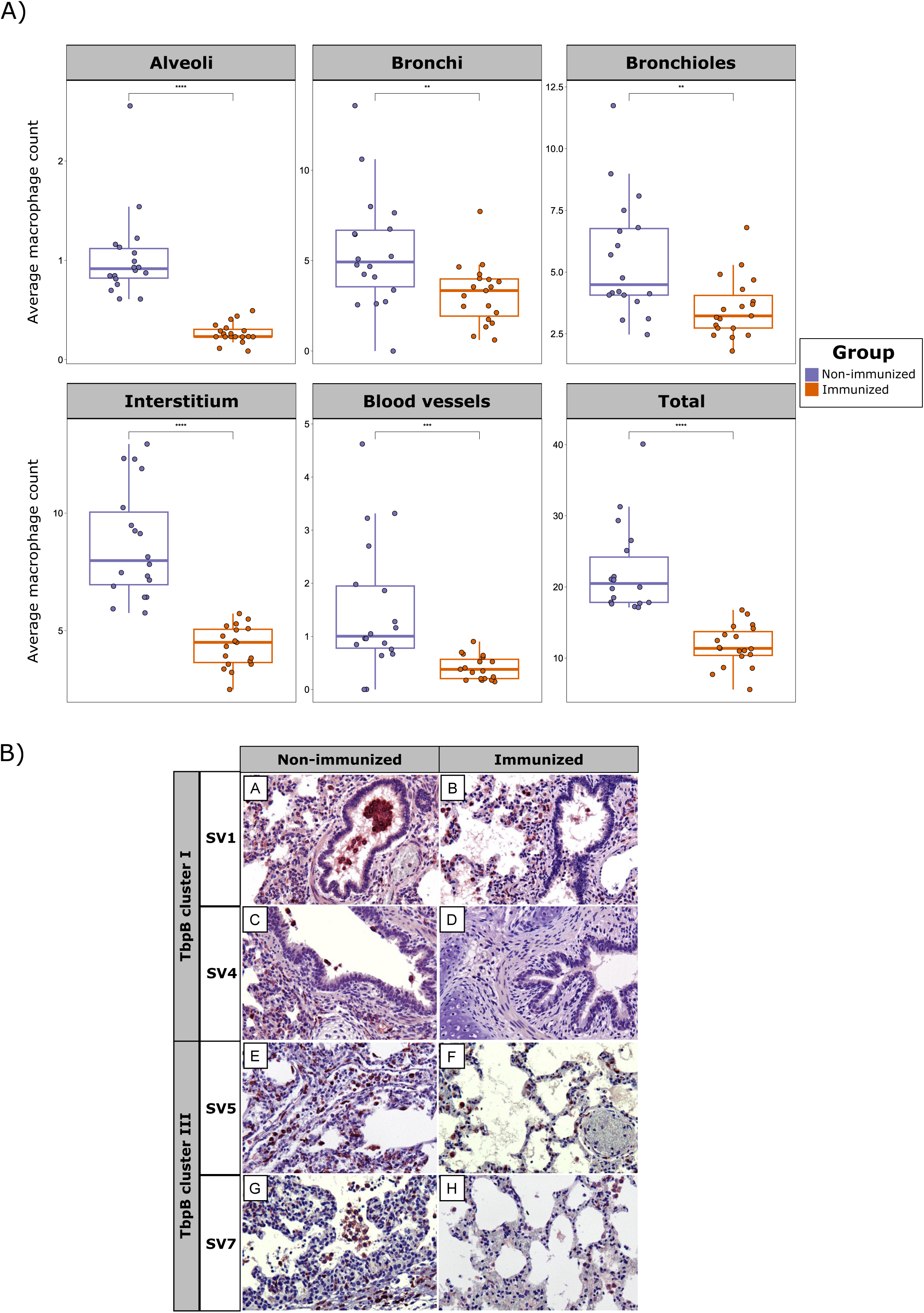
Macrophage abundance in pulmonary tissue after infection challenge in immunized and non-immunized piglets with the oral mucosal TbpB-based vaccine. A) Boxplots representing average macrophage count across various lung locations. Each piglet is represented by a dot with horizontal jitter in the boxplot for visibility. The horizontal box lines represent the first quartile, the median, and the third quartile. Whiskers include the range of points within the 1.5 interquartile range. Differences per group were evaluated with the Wilcoxon rank-sum test. B) Immunohistochemical (IHC) observation of brown labeled CD163 positive macrophages across various lung locations, itemized by *G. parasuis* TbpB cluster and serovar (SV). A and B, bronchioles; C and D, bronchi; E and F, blood vessels and connective tissue; G and H, alveoli. Contrast of nuclei with Harris hematoxylin. Avidin-biotin-peroxidase complex. X100.

## Discussion

Glässer’s disease has a substantial economic impact on the pig industry, particularly affecting nursery, and early fattening stages with significant repercussions on mortality, morbidity, and key productive parameters (2). This underscores the critical need for effective disease management strategies. While vaccination stands as a pivotal measure, the efficacy of current commercial vaccines is hindered by their serovar-specificity (8), which offers limited cross-protection against the diverse pathogenic SVs of *G. parasuis*, urging for the inclusion of multiple serotypes in vaccine formulations (19). The development of subunit vaccines, such as TbpB-based vaccines, may overcome these challenges (11,14). This study demonstrates that mucosal administration of the TbpB^Y167A^ vaccine in colostrum-deprived pigs not only induces robust immunogenicity, but also confers heterologous protection against various prevalent serotypes of field isolates of *G. parasuis* from Spanish farms, thus proving its potential as an effective tool for the control of Glässer’s disease in early stages of production.

The humoral immune response plays an essential role in the protection against the disease (20). Mucosal antibodies, especially IgA, opsonize bacteria facilitating their detection and phagocytosis by alveolar macrophages (21), thus modulating pathogenic colonization in the respiratory tract. The concentration of IgA observed in BALF of immunized colostrum-deprived piglets in this study underscore that this subunit vaccine has significant potential for preventing mucosal colonization by *G. parasuis* in the early stages of life, thereby modulating its pathogenic evolution in nursery piglets. Furthermore, the contribution of specific antibodies in serum susceptibility cannot be underestimated, since it has been demonstrated that the systemic humoral response potentially prevents the spread of *G. parasuis* within the host (22). Our findings, evidenced by a marked increase in IgM levels following vaccination and a subsequent rise in IgG levels against both *G. parasuis* and the target antigen, demonstrate its capacity to induce systemic immunogenicity reaction for controlling the progression of the infection. By enhancing both mucosal and systemic humoral responses, our study aimed to fortify the host ability to counter *G. parasuis* invasion effectively.

Maternally derived antibodies may protect piglets from early colonization by *G. parasuis*, thus underlying the importance of vaccinating sows (3). Recent research by Dellagostin *et al.* (4) has shown that sow vaccination with the TbpB^Y167A^ subunit vaccine effectively reduces *G. parasuis* colonization in suckling piglets. However, as this passive immunity diminishes during lactation, the integration of active piglet immunization becomes essential for controlling the infection at weaning (3). While maternal immunity could potentially interfere with piglet vaccination (2), findings from Frandoloso *et al.* (12) indicate that administering a TbpB-based vaccine directly beneath the oral mucosal epithelium using a needle-free device does not impede the development of an acquired immune response in piglets. This vaccination strategy, as implemented in our research, combined with the observed significant immunogenicity in colostrum-deprived piglets, underscores its promising potential in preventing Glässer’s disease in weaning piglets.

This study presents the first evidence of heterologous protection by TbpB-based vaccines against European clinical field isolates of *G. parasuis*. The TbpB^Y167A^ subunit vaccine, derived from TbpB cluster III (11), has established a precedent for TbpB-based vaccines conferring protection across various capsular types within the homologous TbpB cluster III, including strains belonging to SV5 (11) and SV7 (12). Moreover, it has demonstrated cross-protection across TbpB clusters, effectively against *G. parasuis* SV7 from TbpB cluster I (23). Here, we have extended these findings by corroborating the efficacy of the vaccine against a broader selection of four field clinical isolates from Spanish pig farms, spanning different SVs and TbpB clusters. This effect was manifested by an eightfold increase in overall survival chances and a notable reduction in clinical signs and lesions, regardless of the *G. parasuis* TbpB cluster and SV. It is remarkable, for instance, the consistent heterologous protection in piglets challenged with SV4 (TbpB cluster I), which all survived the challenge period. This outcome contrasts with the variable cross-protection observed with inactivated vaccines (19), underscoring the broad-spectrum efficacy of our vaccine.

This vaccine significantly reduced *G. parasuis* colonization in the respiratory tract and bacterial dissemination to target tissues, such as the tarsus and carpus joints, liver, and brain. These results confirm the role of the heterologous vaccine in enhancing mucosal immunogenicity, preventing bacterial colonization, and halting its further progression. This broad protective capacity is particularly critical, given the role of *G. parasuis* not only as a primary agent in systemic Glässer’s disease but also as a secondary pathogen in the porcine respiratory disease complex (PRDC) (24). By reducing morbidity and mortality, the vaccine stands to significantly improve pig production profitability. With an economic impact of PRDC in Europe estimated between €1.70 to €8.90 per nursery pig and €2.30 to €15.35 per fattening pig (25), the economic and health benefits of our findings cannot be overstated. Consequently, this vaccine emerges as a prime candidate for a universal vaccine against Glässer’s disease.

Interestingly, our study unveiled a notable dichotomy between macroscopic and microscopic lesions in immunized and non-immunized piglets following *G. parasuis* challenge. In non-immunized piglets, we observed a higher incidence of macroscopic lesions, indicative of an acute and severe response to infection, contrasted with a notably lower impact in microscopic lesions. This discrepancy likely reflects the limited survival of these animals, who succumbed within the first three dpi, a timeframe which may be insufficient for the development of extensive microscopic tissue damage. IHC findings supported this observation by revealing an increased presence of macrophages in the lungs of non-immunized piglets. This suggests an intense, yet ultimately overwhelmed, innate defense mechanism attempting to control the infection. Indeed, Macedo *et al.* (26) described that virulent strains of *G. parasuis* require prior opsonization with specific antibodies for effective phagocytosis by alveolar macrophages. Conversely, immunized piglets presented fewer macroscopic lesions, indicating effective mitigation in the impact of the infection. However, these animals developed more microscopic lesions over their extended survival period, indicative of a sustained immune response, a phenomenon previously documented (27). This extended survival allowed for a nuanced interaction between the host immune system and *G. parasuis*, leading to a controlled inflammatory response that not only helped control the infection but also promoted tissue repair processes. Correspondingly, a moderated macrophage response in the lung tissue of immunized piglets likely reflected a more balanced and effective immune strategy. This reduction in the need for extensive inflammatory infiltration, coupled with minimized collateral tissue damage, was facilitated by the *G. parasuis* opsonization by post-vaccinal immunoglobulins, leading to effective phagocytosis by alveolar macrophages (28). This scenario underscores the efficacy of the vaccine not only in preventing severe disease and mortality, but also in modulating the immune response to minimize collateral tissue damage, thus offering a comprehensive protective mechanism against *G. parasuis* infection.

## Conclusions

Our research confirms the cross-protection of TbpB^Y167A^ subunits vaccine against different virulent European *G. parasuis* field isolates, a significant step forward in the control of Glässer’s disease. Demonstrating a heterologous efficacy, this vaccine significantly enhances survival rates, diminishes clinical manifestations, and reduces bacterial colonization and dissemination, regardless of the *G. parasuis* serovar or TbpB cluster. Remarkably, it triggers an immune response that effectively controls infection while promoting tissue repair, evidenced by the balanced macrophage activity in immunized piglets. These findings not only evidence the potential of the vaccine to substantially reduce mortality and morbidity associated with Glässer’s disease, but also highlight its capacity to mitigate its economic impact. Consequently, the TbpB^Y167A^ vaccine emerges as an essential tool for improving pig production profitability, positioning it as a leading candidate for a universal vaccine against this widespread disease in swine.

## List of abbreviations

BALF: Bronchoalveolar lavage fluid
BSA: Bovine serum albumin
CI 95%: Confidence interval 95%
DPI: Days post-infection
HE: Hematoxylin-eosin
IHC: Immunohistochemical
MALDI-TOF MS: Matrix-assisted laser desorption-ionization time-of-flight mass spectrometry
NAD: Nicotinamide adenine dinucleotide
OR: Odds ratio
PBS-T: Phosphate-buffered saline
PPLO: Pleuropneumonia-like organisms
PRDC: Porcine respiratory disease complex
SV: Serovar
Tbp: Transferring-binding protein

## Declarations

### Ethics approval and consent to participate

All procedures involving animals were approved by the institutional bioethical committee (Reference Number OEBA-ULE-003-2022) and performed according to European regulations regarding animal welfare and protection of animals used for experimental and other scientific purposes.

### Consent for publication

Not applicable.

### Availability of data and material

Not applicable.

### Authors’ contributions

Study design was performed by SMM, CBGM and MJGI. Experiment was conducted by AGF, RMP, MPR, APJ, EHL and VAF, with support of SMM and MJGI. Laboratory analyses were performed by AGF with support of MPR and RMP. Statistical analyses were performed by OMA. SMM, MJGI, and OMA provided technical and scientific support on the analysis. AGF, OMA and SMM participated in the manuscript writing or contributed to its revision. All authors revised the manuscript and approved the final version.

## Acknowledgments

We acknowledge the excellent technical assistance provided by María Mediavilla and the contribution in some parts of the research by Miguel Fernández Fernández, particularly in the maintenance and care of the animals. We would also like to thank the generous contribution of Javier Domínguez Juncal, from INIA (Madrid, Spain), for his help with the immunohistochemistry study.

## References

1. Oliveira S, Pijoan C. *Haemophilus parasuis:* New trends on diagnosis, epidemiology and control. Vet Microbiol. 2004;99(1):1–12.

2. Costa-Hurtado M, Barba-Vidal E, Maldonado J, Aragon V. Update on Glässer’s disease: How to control the disease under restrictive use of antimicrobials. Vet Microbiol. 2020;242:108595.

3. Cerdà-Cuéllar M, Naranjo JF, Verge A, Nofrarías M, Cortey M, Olvera A, et al. Sow vaccination modulates the colonization of piglets by *Haemophilus parasuis*. Vet Microbiol. 2010 Oct 26;145(3–4):315–20.

4. Dellagostin D, Klein RL, Giacobbo I, Guizzo JA, Dazzi CC, Prigol SR, et al. TbpB^Y167A^-based vaccine is safe in pregnant sows and induces high titers of maternal derived antibodies that reduce *Glaesserella parasuis* colonization in piglets. Vet Microbiol. 2023 Jan 1;276.

5. Pomorska-Mól M, Markowska-Daniel I, Rachubik J, Pejsak Z. Effect of maternal antibodies and pig age on the antibody response after vaccination against Glässers disease. Vet Res Commun. 2011;35(6):337–43.

6. Blanco-Fuertes M, Correa-Fiz F, López-Serrano S, Sibila M, Aragon V. Sow vaccination against virulent *Glaesserella parasuis* shapes the nasal microbiota of their offspring. Sci Rep. 2022;12(1):1–10.

7. Barasuol BM, Guizzo JA, Fegan JE, Martínez-Martínez S, Rodríguez-Ferri EF, Gutiérrez-Martín CB, et al. New insights about functional and cross-reactive properties of antibodies generated against recombinant TbpBs of *Haemophilus parasuis*. Sci Rep. 2017;7(1):1–13.

8. Liu H, Xue Q, Zeng Q, Zhao Z. *Haemophilus parasuis* vaccines. Vet Immunol Immunopathol. 2016;180:53–8.

9. Curran DM, Adamiak PJ, Fegan JE, Qian C, Yu R hua, Schryvers AB. Sequence and structural diversity of transferrin receptors in Gram-negative porcine pathogens. Vaccine. 2015;33(42):5700–7.

10. Martín de la Fuente AJ, Gutiérrez Martín CB, Pérez Martínez C, García Iglesias MJ, Tejerina F, Rodríguez Ferri EF. Effect of Different Vaccine Formulations on the Development of Glässer’s Disease Induced in Pigs by Experimental *Haemophilus parasuis* Infection. J Comp Pathol. 2009 Feb;140(2–3):169–76.

11. Frandoloso R, Martínez-Martínez S, Calmettes C, Fegan J, Costa E, Curran D, et al. Nonbinding site-directed mutants of transferrin binding protein B exhibit enhanced immunogenicity and protective capabilities. Infect Immun. 2015;83(3):1030–8.

12. Frandoloso R, Chaudhuri S, Frandoloso GCP, Yu RH, Schryvers AB. Proof of Concept for Prevention of Natural Colonization by Oral Needle-Free Administration of a Microparticle Vaccine. Front Immunol. 2020 Oct 23;11.

13. Fernández AG, Martín CBG, Rilo MP, Fernández EP, Pérez RM, Frandoloso R, et al. Phylogenetic study and comparison of different TbpB obtained from *Glaesserella parasuis* present in Spanish clinical isolates. Res Vet Sci. 2023 Apr 1;157:35–9.

14. Guizzo JA, Chaudhuri S, Prigol SR, Yu RH, Dazzi CC, Balbinott N, et al. The amino acid selected for generating mutant TbpB antigens defective in binding transferrin can compromise the in vivo protective capacity. Sci Rep. 2018 Dec 1;8(1).

15. Domenech N, Rodríguez-Carreño MP, Filgueira P, Alvarez B, Chamorro S, Domínguez J. Identification of porcine macrophages with monoclonal antibodies in formalin-fixed, paraffin-embedded tissues. Vet Immunol Immunopathol. 2003;94(1–2):77–81.

16. R Core Team. R: A Language and Environment for Statistical Computing. Vienna, Austria: Foundation for Statistical Computing; 2023.

17. Benjamini Y, Hochberg Y. Controlling the False Discovery Rate: A Practical and Powerful Approach to Multiple Testing. R Stat Soc. 1995.

18. Wickham H. ggplot2: Elegant Graphics for Data Analysis. Springer-Verlag New York. 2016.

19. Zhao Z, Liu H, Xue Y, Chen K, Liu Z, Xue Q, et al. Analysis of efficacy obtained with a trivalent inactivated *Haemophilus parasuis* serovars 4, 5, and 12 vaccine and commercial vaccines against Glässer’s disease in piglets. Can J Vet Res. 2017;81(1):22–7.

20. Nedbalcova K, Kucerova Z, Krejci J, Tesarik R, Gopfert E, Kummer V, et al. Passive immunisation of post-weaned piglets using hyperimmune serum against experimental *Haemophilus parasuis* infection. Res Vet Sci. 2011;91(2):225–9.

21. Olvera A, Ballester M, Nofrarias M, Sibila M, Aragon V. Differences in phagocytosis susceptibility in *Haemophilus parasui*s strains. Vet Res. 2009;40(3).

22. Brockmeier SL, Loving CL, Mullins MA, Register KB, Nicholson TL, Wiseman BS, et al. Virulence, transmission, and heterologous protection of four isolates of *Haemophilus parasuis*. Clin Vaccine Immunol. 2013;20(9):1466–72.

23. Prigol SR, Klein R, Chaudhuri S, Frandoloso GP, Guizzo JA, Martín CBG, et al. TbpB^Y167A^-Based Vaccine Can Protect Pigs against Glässer’s Disease Triggered by *Glaesserella parasuis* SV7 Expressing TbpB Cluster I. Pathogens. 2022;11(7).

24. Schuwerk L, Hoeltig D, Waldmann KH, Strutzberg-Minder K, Valentin-Weigand P, Rohde J. Serotyping and pathotyping of *Glaesserella parasuis* isolated 2012–2019 in Germany comparing different PCR-based methods. Vet Res. 2020;51(1):1–14.

25. Boeters M, Garcia-Morante B, van Schaik G, Segalés J, Rushton J, Steeneveld W. The economic impact of endemic respiratory disease in pigs and related interventions - a systematic review. Porc Heal Manag. 2023;9(1).

26. Macedo N, Rovira A, Torremorell M. *Haemophilus parasuis*: Infection, immunity and enrofloxacin. Vet Res. 2015;46(1):1–6.

27. Frandoloso R, Martínez S, Rodríguez-Ferri EF, García-Iglesias MJ, Pérez-Martínez C, Martínez-Fernández B, et al. Development and characterization of protective *Haemophilus parasuis* subunit vaccines based on native proteins with affinity to porcine transferrin and comparison with other subunit and commercial vaccines. Clin Vaccine Immunol. 2011;18(1):50–8.

28. Matiašková K, Kavanová L, Kulich P, Gebauer J, Nedbalcová K, Kudláčková H, et al. The Role of Antibodies Against the Crude Capsular Extract in the Immune Response of Porcine Alveolar Macrophages to In Vitro Infection of Various Serovars of *Glaesserella (Haemophilus) parasuis*. Front Immunol. 2021;12(April):1–15.

